# MetaPGN: a pipeline for construction and graphical visualization of annotated pangenome networks

**DOI:** 10.1101/398636

## Abstract

Pangenome analyses facilitate the interpretation of genetic diversity and evolutionary history of a taxon. However, there is an urgent and unmet need to develop new tools for advanced pangenome construction and visualization, especially for metagenomic data. Here we present an integrated pipeline, named MetaPGN, for construction and graphical visualization of pangenome network from either microbial genomes or metagenomes. Given either isolated genomes or metagenomic assemblies coupled with a reference genome of the targeted taxon, MetaPGN generates a pangenome in a topological network, consisting of genes (nodes) and gene-gene genomic adjacencies (edges) of which biological information can be easily updated and retrieved. MetaPGN also includes a self-developed Cytoscape plugin for layout of and interaction with the resulting pangenome network, providing an intuitive and interactive interface for full exploration of genetic diversity. We demonstrate the utility of MetaPGN by constructing *Escherichia coli* (*E. coli*) pangenome networks from five *E. coli* pathogenic strains and 760 human gut microbiomes respectively, revealing extensive genetic diversity of *E. coli* within both isolates and gut microbial populations. With the ability to extract and visualize gene contents and gene-gene physical adjacencies of a specific taxon from large-scale metagenomic data, MetaPGN provides advantages in expanding pangenome analysis to uncultured microbial taxa. MetaPGN is available at https://github.com/peng-ye/MetaPGN.

## Introduction

The concept of the pangenome, defined as the full complement of genes in a clade, was first introduced by Tettelin *et al.* in 2005 [1]. Pangenome analyses of a species now provide insights into core-and accessory-genome profiles, within-species genetic diversity, evolutionary dynamics and niche-specific adaptions. A number of methods and tools have to date been proposed for pangenome analysis on genomic or metagenomic data (Table 1).

**Table 1.**
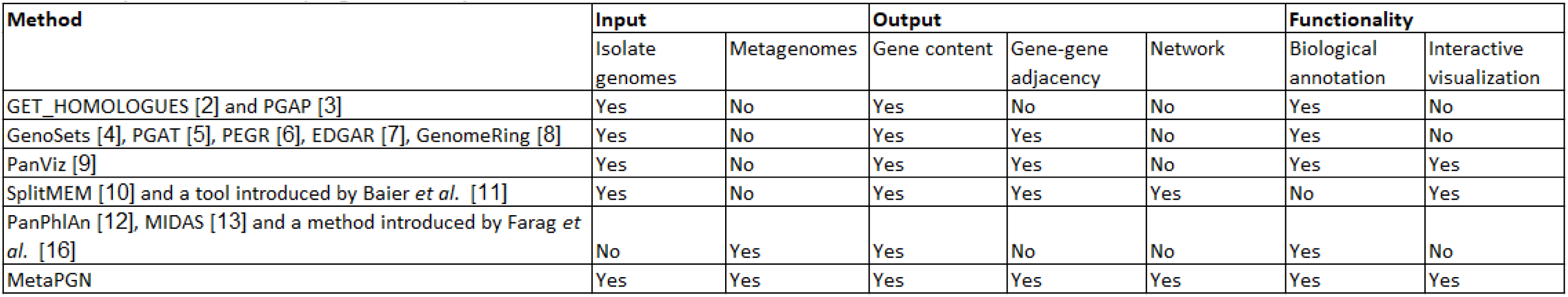
Comparison of serveral pangenome analysis methods.

Typical pangenome tools such as GET_HOMOLOGUES [2] and PGAP [3], mainly focus on analyzing homologous gene families and calculating the core/accessory genes of a given taxon. However, these tools cannot provide the variations of gene-gene physical relationships. Tools like GenoSets [4], PGAT [5], PEGR [6], EDGAR [7], GenomeRing [8] and PanViz [9] are developed to generate a linear or circular presentation of compared genomes, which can indicate the physical relationships between genomic sequences or genes. However, in the linear or circular representations generated by these tools, the same homologous region is visualized multiple times and shown on separate input genomes. Hence, it will be difficult for users to track a homologous region among the input genomes, especially when there is a large number of homologous regions and input genomes.

Pangenomes built using *de Bruijn* graph, like SplitMEM [10] and a tool introduced by Baier *et al.* [11], partly solve the above problems. In the resulting graph generated by these tools, the complete pangenome is represented in a compact graphical representation such that the core/accessory status of any genomic sequences is immediately identifiable, along with the context of the flanking sequences. This strategy enables powerful topological analysis of the pangenome not possible from a linear/circular representation. Nevertheless, tools based on the *de Bruijn* graph algorithm can only construct a compact network comprised of core/accessory genomic sequences instead of genes, which means retrieving or updating functional information in downstream analysis will be difficult. Furthermore, these tools do not visualize the constructed *de Bruijn* graph and provide an interactive interface for users to explore the graph.

Moreover, all the above-mentioned tools analyze pangenomes via genomic data which require organisms isolated from the environment and cultured *in vitro*. Recent advances in metagenomics have led to a paradigm shift in pangenome studies from a limited quantity of cultured microbial genomes to large-scale metagenomic datasets containing huge potential for functional and phylogenetic resolution from the still uncultured taxa. Several existing tools dealing with metagenomic data are based on constructed pangenomes and cannot utilize the abundant gene resources contained in metagenomes to extend the pangenomes in question. For example, PanPhlAn [12], MIDAS [13], and a pipeline introduced by Delmont and Eren [14] maps reads onto a reference pangenome, to describe the pattern of the presence/absence of genes in metagenomes. As for another example, Kim *et al.* [15] clustered genes predicted from metagenomic contigs with *Bacillus* core genes for profiling the *Bacillus* species in the microbiomes. Recently, Farag *et al.* [16] aligned metagenome contigs with reference genomes for identification of *“Latescibacteria”* genomic fragments. Even though this strategy can theoretically recruit sequences not present in the reference genomes, it is likely to filter out *“Latescibacteria”* genomic fragments with structural variations compared to the reference ones. Furthermore, all these aforementioned methods using metagenomic data do not organize the pangenome using a network, which is essential for efficiently storage and visualization of pangenomes constructed from metagenomic data.

Here, we introduce an integrated pipeline (MetaPGN) for network-based construction and visualization of prokaryotic pangenomes for both isolated genomes and metagenomes. Given genomic or metagenomic assemblies and a reference genome of a taxon of interest, MetaPGN derives a pangenome network for integrating genes (nodes) and gene-gene adjacencies (edges) belonging to a given taxon. MetaPGN also includes a specific Cytoscape plugin for layout of and interaction with the resulting pangenome network, providing an intuitive and interactive interface for the exploration of gene diversity. For example, in the visualized network in Cytoscape, users can specify gene annotations, customize the appearance of nodes and edges, and search and concentrate on genes of certain functions. We applied MetaPGN on assemblies from five pathogenic *E. coli* strains and 760 human gut microbiomes respectively, with *E. coli* K-12 substr. MG1655 (*E. coli* K-12) being the reference genome. Our results showed that by taking gene adjacency into account and visualizing the pangenome network in a well-organized manner, MetaPGN can assist in illustrating genetic diversity in genomic or metagenomic assemblies graphically and conveniently.

## Results

### General workflow

MetaPGN accepts genome or metagenome assemblies as input (query assemblies) and requires a reference genome for recruitment of the query assemblies and as the skeleton of the pangenome network. The MetaPGN pipeline can be divided into two main parts: (i) construction of a pangenome network comprised of representative genes, including gene prediction, gene redundancy elimination, gene type determination, pairwise gene adjacency extraction, assembly recruitment (for metagenomic assemblies), and pangenome network generation, and (ii) visualization of the pangenome network in an organized way, where nodes represent genes and edges indicate gene adjacencies, in Cytoscape [17] with a self-developed plugin (Fig. 1, Fig. S1, and Methods). From the resultant pangenome network, the degree of similarity among homologous genes, as well as their genomic context is easily visible. Of note, users can further add and update annotation for nodes and edges in the networks, based on which elements of interest can be accessed conveniently.

**Figure 1.**
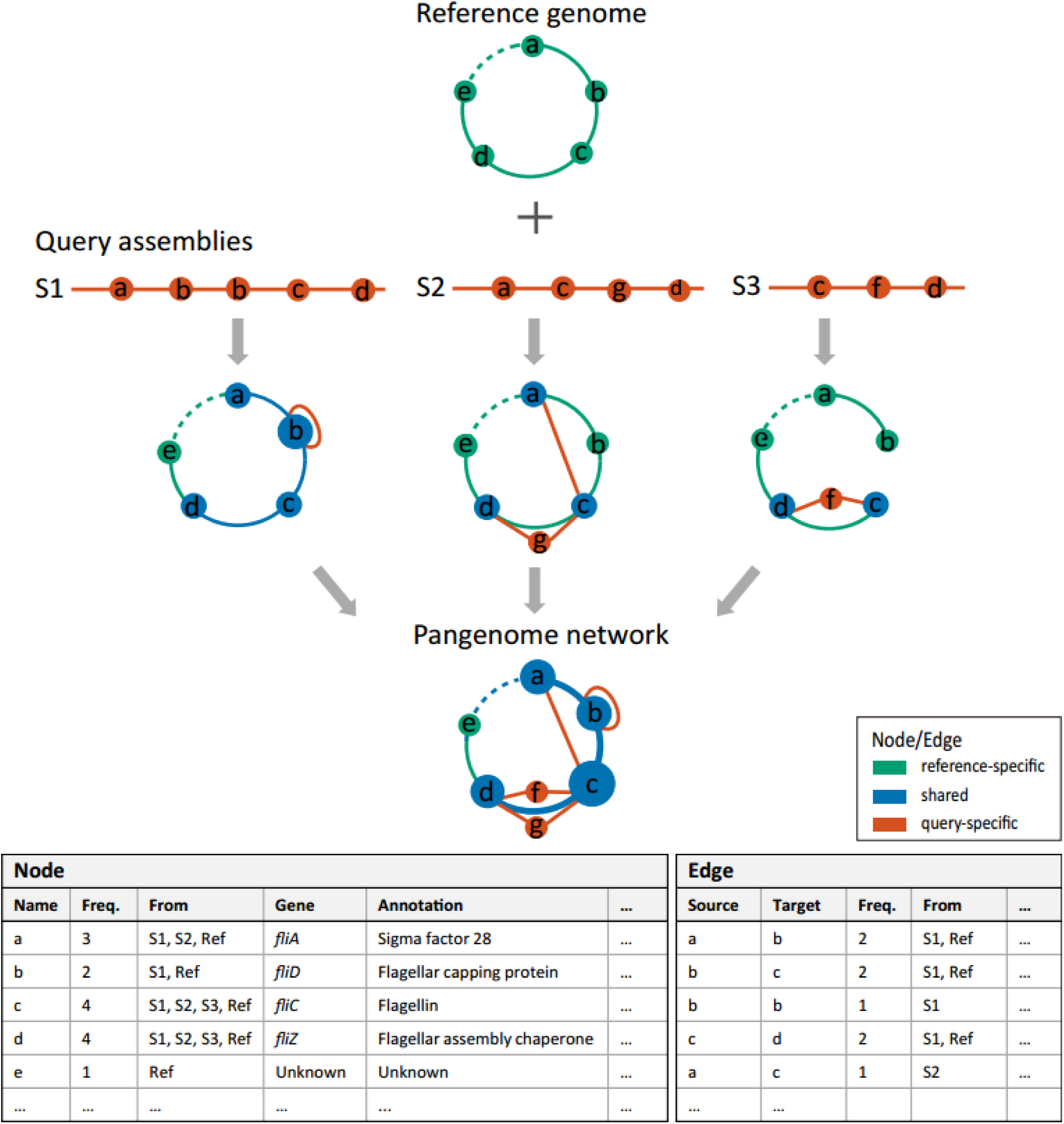
An Overview of the MetaPGN pipeline: from assemblies to a pangenome network. Gene prediction is performed on query assemblies. The resulting genes are clustered, after which genes in the same cluster are represented by the longest sequence of this cluster called the representative gene (node a-g). All these representative genes are then aligned against genes on the given reference genome. From the alignment result, genes shared between the representative gene set and the reference gene set are defined as ‘shared’ genes (blue). The remaining representative and reference genes other than those shared genes are defined as ‘query-specific’ genes (red) and ‘reference-specific’ genes (green), respectively. Pairwise gene physical adjacency of representative genes on the query assemblies and of reference genes are then extracted, and status for each adjacency of being ‘shared’ (blue), ‘query-specific’ (red), or ‘reference-specific’ (green) is determined. Finally, based on the recruited assemblies and the reference genome, a pangenome network is generated: each node stands for a reference gene or a representative gene on the recruited assemblies; two nodes are connected by an edge if they are physically adjacent on the recruited assemblies or the reference genome. The weight of a node or an edge is its occurrence frequency on all of the recruited assemblies and the reference genome (Methods). The pangenome network is then visualized in Cytoscape with a self-developed plugin (Methods) for a better arrangement. Biological information of nodes and edges, such as gene name and annotation, can be easily retrieved in the interactive user interface in Cytoscape.

### Pangenome network of 5 pathogenic *Escherichia coli* genomes

In order to demonstrate its potential in studying microbial genetic diversity and phenotype-genotype relationship, we first applied MetaPGN on genomes of 5 pathogenic *E. coli* isolates, *E. coli* O26:H11 str. 11368, *E. coli* O127:H6 E2348/69, *E. coli* O157:H7 str. EDL933, *E. coli* O104:H4 str. 2011C-3493 and *E. coli* 55989. A commensal *E. coli* strain, K-12 substr. MG1655 (Supplementary Table S1) was chosen as the reference genome in this instance and in all examples shown below.

A pangenome network consisting of 9,161 nodes and 11,788 edges (Supplementary Table S3, Supplementary File 2) was constructed and visualized (Methods). Based on the well visualized pangenome network along with functional annotation, we can now graphically observe the extent of variations of certain genes, as well as their genomic context. For example, when focusing on a cluster of flagellar genes (Fig. 2a), we found that *fliC* sequences encoding the filament structural protein (H-antigen) and *fliD* sequences encoding the filament capping protein are highly divergent with nucleotide sequence identity < 95% and/or overlap < 90% among these *E. coli* strains (See Methods). In contrast, four genes encoding chaperones (*fliS, fliT, fliY, fliZ*) and a gene related to regulation of expression of flagellar components (*fliA*) are conserved (nucleotide sequence identity ≥95% and overlap ≥ 90%) over all the *E. coli* strains investigated. A gene (270bp) encoding a hypothetical protein is uniquely presented between *fliC* and *fliD* in *E. coli* O157:H7 str. EDL933.

**Figure 2.**
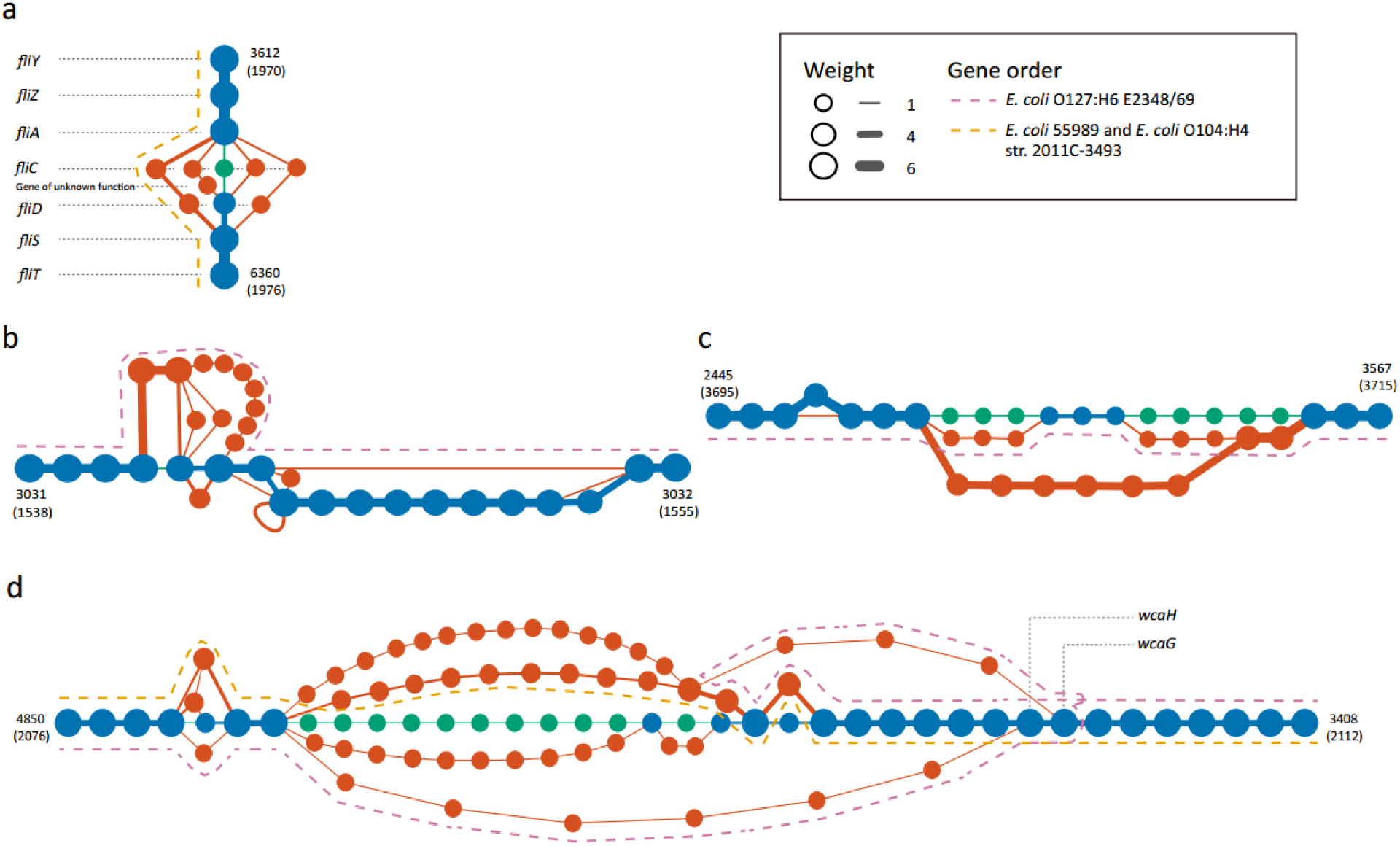
Subgraphs of highly variable genes in the pangenome network of 5 pathogenic *E. coli* strains (manually arranged). (a) a cluster of flagellar genes. (b) a cluster containing outer membrane protein-coding genes. (c) a cluster of genes responsible for biosynthesis of the O antigen. (d) another cluster of O antigen-related genes. Green, blue, red nodes and edges denote reference-specific, shared, and query-specific genes and gene adjacencies, respectively. Size of nodes and thickness of edges indicates their weight (occurrence frequency). Numbers alongside shared genes are their indexes in the representative gene set.

In a fimbria protein-related gene cluster, compared to the reference *E. coli* strain, all the 5 pathogenic strains possess several genes located between two conserved genes encoding an outer membrane protein and a regulatory protein, and *E. coli* O127:H6 E2348/69 uniquely exhibits more genes encoding proteins of unknown functions (Fig. 2b).

For a gene cluster responsible for the biosynthesis of lipopolysaccharides (LPS), *E. coli* O127:H6 E2348/69 shares three genes with the reference strain that differentiate from the other 4 pathogenic strains (Fig. 2c). For another gene cluster of related function, the *E. coli* O127:H6 E2348/69 also shows a strain-specific duplication event of two genes involved in colanic acid (CA) synthesis (wcaH and wcaG, denoted by a purple dash line in Fig. 2d). It has been demonstrated that CA can modify lipopolysaccharide (LPS) generating a novel form (MLPS) which may enhance survival of *E. coli* in different ways [18]. The two wcaH genes in *E. coli* O127:H6 E2348/69, may even though they share high similarity (99.1% identity) confer the strain with different functional potentials for CA formation and thereby novel survival mechanisms.

In addition, the German outbreak *E. coli* O104:H4 str. 2011C-3493 shares identical nodes and edges in the flagellar-related gene cluster (Fig. 2a) and the O antigen-related gene cluster with a historical *E. coli* 55989 (Fig. 2d), suggesting a close evolutionary relationship between these strains as previously reported [19,20].

These results demonstrate the feasibility of MetaPGN for construction and visualization of microbial pangenomes in an organized way. Moreover, by involving genomic adjacency and offering easy-to-achieve biological information, MetaPGN provides a convenient way to assist biologists in exposing genetic diversity for genes of interest among the organisms under study.

### Pangenome network of *E. coli* in 760 metagenomes

Moving beyond surveying the pangenome network of isolate genomes, we applied MetaPGN in metagenomic datasets to interrogate the *E. coli* pangenome network on a grander scale. Assemblies of 760 metagenomes sequenced in the Metagenomics of the Human Intestinal Tract (MetaHIT) project [21–24] were collected, which contained 8,096,991 non-redundant genes with annotations [24]. As metagenome assemblies are from varied taxa, it is necessary to recruit assemblies of the targeted taxon before construction of the pangenome network. In this study, metagenome assemblies were recruited using a gene alignment-based strategy, which was assessed with mock datasets (Methods). With the recruited assemblies, a pangenome network consisting of 9,406 nodes and 14,676 edges (Supplementary Table S3, Supplementary File S3) was generated and visualized after refinement (Methods).

Based on annotation, we first searched flagellin-related genes in this network. We found that the pattern of adjacencies among these genes was similar to that in the pangenome network of the 5 pathogenic *E. coli* genomes: *fliC* and *fliD* are hypervariable while *fliT, fliY, fliZ* and *fliA* are very conserved among these 760 samples. However, some genes of unknown function locate between *fliC* and *fliA* (Fig. 3a), instead of between *fliC* and *fliD* in the pangenome network of the 5 pathogenic *E. coli* strains (Fig 2a).

**Figure 3.**
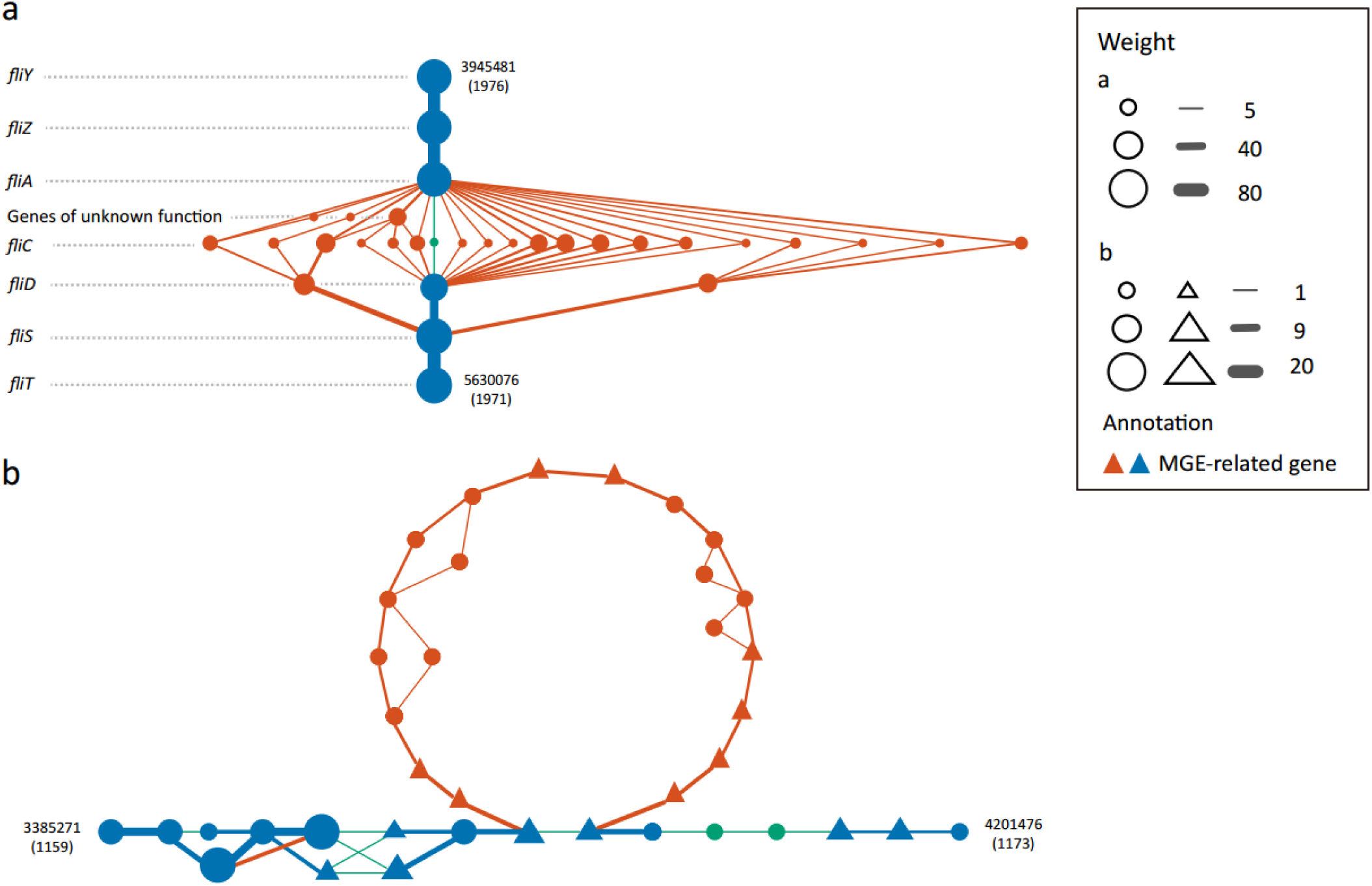
Two subgraphs of the pangenome network of *E. coli* constructed from 760 metagenomes (manually arranged). (a) a cluster of flagellar genes. (b) a cluster of genes containing MGEs. Green, blue, red nodes and edges denote reference-specific, shared, and query-specific genes and gene adjacencies. Triangles represent MGEs. Size of nodes and thickness of edges indicates their weight (occurrence frequency). Numbers alongside shared genes are their indexes in the representative gene set.

We then investigated mobile genetic elements (MGEs) in this pangenome network, as they can induce various types of genomic rearrangements[25]. Of the 362 nodes (∼4%) annotated as MGE-related (according to Cluster of Orthologous Groups annotation done in reference [24]), many were flanked by shared genes on different *E. coli* genomes. In a region of the network, a gene cluster containing MGEs is query-specific, indicating there might be genomic rearrangements caused by strain-specific MGEs within the *E. coli* species (Fig. 3b). In another part of the network harboring MGEs, we observed that several branches of non-MGE genes are inserted in between two MGEs, which may imply a mutation hotspot within the region, or the existence of MGEs as yet undescribed (Fig. S1).

Application of MetaPGN in large-scale metagenomic data generated an *E. coli* pangenome network that might hardly be constructed from isolated genomes. As demonstrated here, the assembly-recruitment based, well-organized and visualized pangenome network can greatly expand our understanding in the genetic diversity of a taxon, although further efforts in bioinformatic and experimental analyses are needed to verify and extend these findings.

### Assessment of pangenome networks derived from metagenomes

Affected by the complexity of microbial communities, limitations in sequencing platforms and imperfections of bioinformatic algorithms, a genomic sequence of an organism is frequently split into dozens of assemblies when assembled from metagenomic reads. Due to this nature, a pangenome network recovered from a limited number of assemblies is likely to be segmented compared to a complete genome. To propose a minimum size of assemblies for getting an approximately complete connected pangenome network, we assessed the completeness of *E. coli* pangenome networks derived from varying size of recruited assemblies (Methods). As shown in Fig. 4, the count of connected subnetworks drops dramatically with the total length of recruited assemblies increasing from 5 Mb to 50 Mb (roughly from 1 × to 10 × of a *E. coli* genome), then barely changes even when using all recruited assemblies of the dataset (215 Mb, from 760 samples). Based on this analysis, a minimum size of recruited assemblies 10-fold of the studied genome is required to generate a relatively intact pangenome network when constructed from metagenomes.

**Figure 4.**
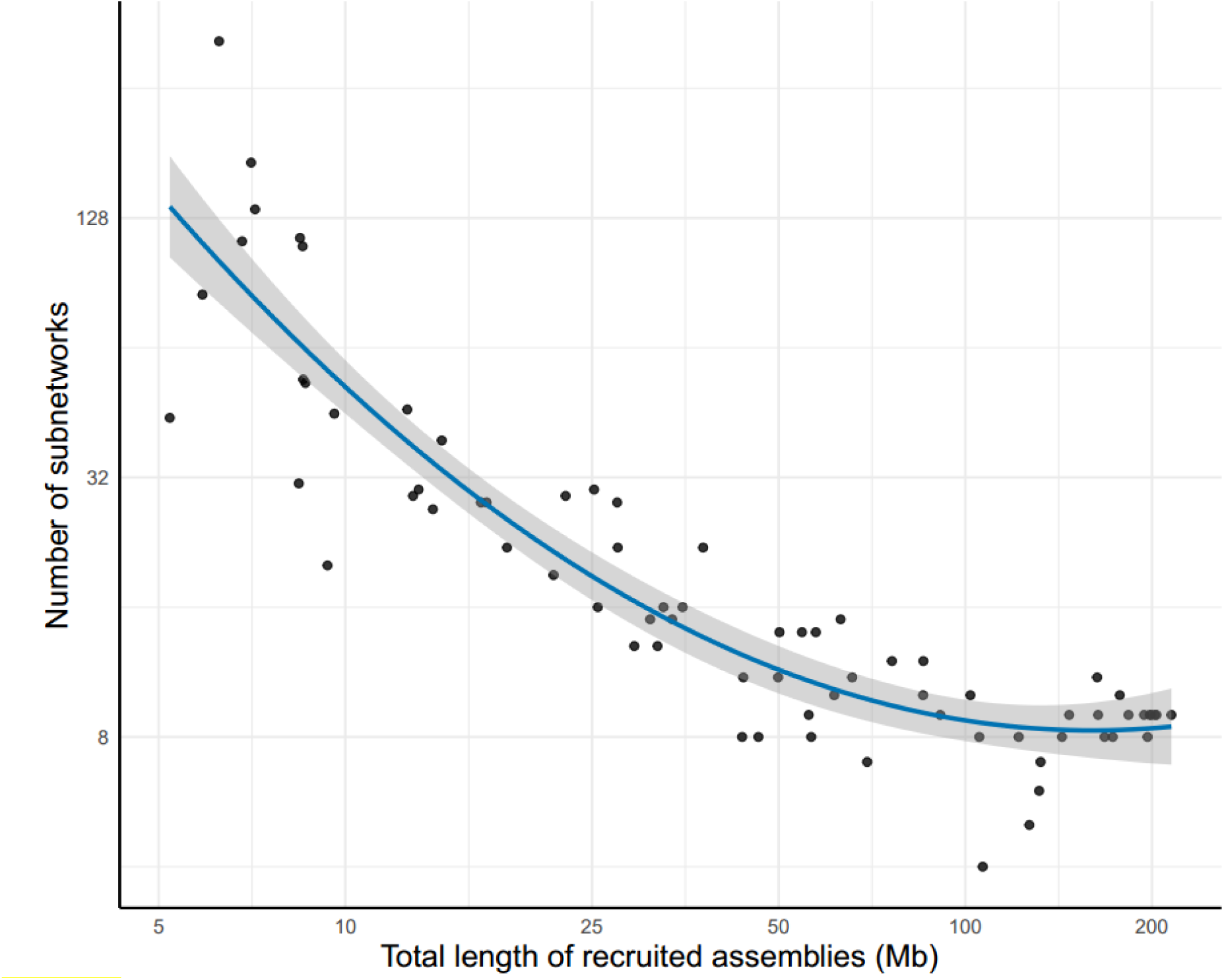
Number of subnetworks in pangenome networks derived from varying sizes of recruited assemblies. The x-axis indicates total length of recruited assemblies for each sub-dataset and the y-axis represents the number of subnetworks in the pangenome network derived from each sub-dataset. The curve was fitted for the scatters using the ‘loess’ smoothing method in R[47]. The shaded area displays the 95% confidential intervals of the curve. Axes are log2-transformed.

## Discussion

Since first coined more than a decade ago, pangenome analysis has provided a framework for studying the genomic diversity within a species. Current methods for pangenome analyses mainly focus on gene contents but ignore their genomic context, as well as having shortages in pangenome visualization. Besides, available methods are usually designed for genomic data and not capable of constructing pangenomes from metagenomics data. To fill these gaps, our MetaPGN pipeline takes genome or metagenome assemblies as input, uses gene contents as well as pairwise gene adjacency to generate a compact graphical representation for the gene network based on a reference genome, and visualizes the network in Cytoscape with a self-developed plugin (Fig. 1, Fig. S2).

From the two MetaPGN-derived *E. coli* pangenome networks, we can directly observe the diversity of genes among the five pathogenic *E. coli* strains and 760 human gut microbiomes with respect to the reference genome. For instance, in the pangenome network for the 5 pathogenic *E. coli* strains, we found that nucleotide sequences of the *fliC* gene which carries H-antigen specificity were highly divergent among the *E. coli* assemblies (Fig. 2a). These *fliC* sequences were more varied in the 760 human gut microbiomes (Fig. 3a). In addition, genes for synthesis of O-antigen and outer membrane protein showed a great diversity in the pangenome network of the 5 E. coli strains (Fig. 2c, Fig. 2d). These results are in agreement with previous findings on H-antigen specificity related genes [26–28] and O-antigen related genes [29,30]. We also showed that when gene adjacency is incorporated into the construction and visualization of pangenomes, locations of genes of unknown function are identified, which may be helpful for the inference of their biological functions. For example, in both the two pangenome networks, we found genes of unknown function locating between the *fliC* gene and other flagellin-related genes (Fig. 2a, located between *fliC* and *fliD*, Fig. 3a, and located between *fliC* and *fliA*), indicating that these functionally unknown genes may play a role in flagellin biosynthesis [31], although further experimental trials are needed to prove this point. Additionally, from the pangenome network of the five *E. coli* strains, we observed a variation in *E. coli* O127:H6 E2348/69, which was shown to stem from a duplication event of two genes involved in colanic acid synthesis (*wcaH* and *wcaG*, Fig 2d). This finding indicates that knowledge of genomic adjacency may also shed light on structural variations among the input assemblies. If extended, genomic adjacency may further help in finding possible functional sequences which are associated with structural variations, as Delihas [32] and Wang *et al.* [33] reported on repeat sequences concentrated at the breakpoints of structural variations. Studying genomic adjacency can also improve the discovery of potential functional modules, as Doron *et al.* [34] systematically discovered bacterial defensive systems by examining gene families enriched next to known defense genes in prokaryotic genomes. These examples illustrate the value of including gene adjacencies in visualizing a pangenome to retrieve biological information. Although the examples shown in this study use the genome of a commensal *E. coli* strain for assembly recruitment and network arrangement, users can specify the reference genome when applying MetaPGN. Epidemiologists can use MetaPGN to compare assemblies of outbreak strains or viruses, such as *Vibrio cholerae* or Ebola virus, with those of some well-studied pathogenic strains to find novel variations involved in pathogenesis, which may further provide candidate targets for drug and vaccine design [35,36].

Genomic variants of intestinal bacteria were found to be correlated with diseases. As one example, among the common members of the normal colonic microbiota, *Bacteroides fragilis* (*B. fragilis*), the inclusion of a pathogenicity island (BfPAI) distinguished enterotoxigenic strains (ETBF) from nontoxigenic ones (NTBF), by their ability to secrete a zinc-dependent metalloprotease toxin that can induce inflammatory diarrhea and even colon carcinogenesis [37,38]. As another example, Scher *et al.* performed shotgun sequencing on fecal samples from newly-onset untreated rheumatoid arthritis (NORA) patients and healthy individuals, and identified several NORA-specific *Prevotella copri* genes [39]. Hence, pangenome networks built from metagenomes of patients and healthy subjects may aid in detecting associated or causal genomic variants of a certain species.

It should be noticed that, in this pipeline, we compare genes depending on nucleotide-level sequence identity and overlap: genes with ≥ 95% identity and ≥ 90% overlap are regarded as the same gene. However, genes sharing the same function may not satisfy this criterion (≥95% identity and ≥90% overlap), and protein encoded by these genes may exhibit more similarity due to different codon usage. Hence, in our future work, we intend to cluster genes by comparing their nucleotide sequences as well as the amino acid sequences. Furthermore, the current MetaPGN pipeline does not consider other genomic features or physical distances between genes in constructing the pangenome network. Thus, differences in other genomic features such as ribosomal binding site (RBS) sequences [40,41] and distances between the RBS and start codons [42] may result in distinct phenotypes. Accordingly, users may include such information in analyzing pangenome networks.

To conclude, MetaPGN enables direct illustration of genetic diversity of a species in pangenome networks, improving understanding of genotype-phenotype relationships and evolutionary history.

## Methods

### Pangenome network construction in MetaPGN

First, gene prediction of query assemblies is performed using MetaGeneMark (Version 2.8) [43]. In order to eliminate redundancy, the resultant genes are clustered by CD-HIT (Version 4.5.7) [44] with identity ≥95% and overlap ≥90, and genes in a same cluster are represented by the longest sequence of the cluster which is termed the representative gene. Representative genes of all clusters are subsequently aligned against genes on the given reference genome using BLAT (Version 34) [45]. From the alignment result, genes shared between the representative gene set and the reference gene set with identity ≥95% and overlap ≥90% are defined as ‘shared genes’. The remaining representative and reference genes other than those shared genes are defined as ‘query-specific genes’ and ‘reference-specific genes’, respectively. Pairwise gene physical adjacency of representative genes on the query assemblies and of reference genes are then extracted, and status for each adjacency of being ‘shared’, ‘query-specific’, or ‘reference-specific’ is determined. Finally, based on the recruited assemblies and the reference genome, an initial pangenome network is generated: each node stands for a reference gene or a representative gene on the recruited assemblies; two nodes are connected by an edge if they are physically adjacent on the recruited assemblies or on the reference genome. The weight of a node or an edge denotes its occurrence frequency on all of the recruited assemblies and the reference genome.

### Pangenome network visualization in MetaPGN

The following preprocessing work on the initial pangenome network was implemented before visualization: 1. The initial pangenome network was refined by removing isolated networks (networks not connected with the backbone) and tips (nodes only connected with another node); 2. Nodes and edges were added with some extra attributes, such as the status of the nodes and edges (query-specific, reference-specific or shared), whether the genes for the nodes were phage-, plasmid-, CRISPR-related genes and so on (Supplementary Table S3). Users can specify the attributes of nodes and edges according to their own datasets.

We then used a self-developed Cytoscape plugin to visualize the pangenome network in an organized way (Supplementary Text 2 in Supplementary File S1 illustrates how to install and use the plugin in Cytoscape). Our algorithm for organizing nodes in the network is as follows:

1. Construct a circular skeleton for the pangenome network with shared nodes and reference-specific nodes, according to positions of their related reference genes on the reference genome. If there are two or more representative genes similar to the same reference gene (≥95% identity and ≥90% overlap), use one of these representative genes to construct the skeleton and place the others on both sides of the skeleton in turn (Fig. S2 a).
2. Arrange query-specific nodes region by region, including, 2.1. Select query-specific nodes in a region spanning less than 30 nodes in the skeleton (see Supplementary Text 3 in Supplementary File S2 for more details). 2.2. Arrange these query-specific nodes as follows,
  i. For those that directly link with two nodes on the skeleton, place them on the bisector of the two skeleton nodes. If there are two or more query-specific nodes directly linking with the same pair of nodes on the skeleton, place them on both sides of the bisector of these pair of skeleton nodes in turn (Fig. S2 b).
  ii. Among the remaining nodes, for those that directly link with two placed nodes, place them on the bisectors of the placed ones. Iterate this step for five times (Fig. S2 c).
  iii. For the remaining nodes, place them into an arc without moving the placed nodes (Fig. S2 d), or else place them one by one starting near a placed node (Fig. S2 e).

### Construction and visualization of the 5-*E. coli*-genome pangenome network

Genes were extracted from the complete genome for each strain (Supplementary Table S1). With *E. coli* K-12 as the reference, a pangenome network was generated for these five *E. coli* strains using our MetaPGN tool. In the visualization of this pangenome network, we used green, blue and red color to denote a reference-specific, shared, and query-specific node or edge, respectively, and specified sizes of nodes and widths of edges with their occurrence frequency in the input genomes.

### Assessment of the gene alignment-based assembly recruitment strategy

A gene alignment-based strategy was used for recruitment of metagenome assemblies in this study, which considers 1) the count of genes on an assembly (*c*), and 2) the ratio of the number of shared genes (designated as aforementioned) on an assembly to the total number of genes on that assembly (*r*). *c* = 3 paired with *r* = 0.5, requiring at least 3 genes including 2 shared genes containing in an assembly, was chosen for recruitment of metagenome assemblies in this study.

5 mock metagenomic datasets were used to assess the performance of this strategy. Briefly, simulated reads of 60 bacterial genomes from 14 common genera (*Bifidobacterium, Clostridium, Enterobacter, Escherichia, Haemophilus, Klebsiella, Lactobacillus, Neisseria, Pseudomonas, Salmonella, Shigella, Staphylococcus, Streptococcus, Yersinia*) present in the human gut (Supplementary Table S1), including the 5 pathogenic *E. coli* strains mentioned above and 10 strains from *E. coli*-closely-related species (*Enterobacter aerogenes, Enterobacter cloacae, Escherichia albertii*, Escherichia *fergusonii, Klebsiella oxytoca, Klebsiella pneumoniae, Shigella boydii, Shigella sonnei* and *Salmonella enterica*), were generated by iMESSi [46]. Each dataset was simulated at the same complexity level with 100 million (M) 80bp paired-end reads of 12 strains from 11-12 different genera, including 2 strains of closely related species to *E. coli*, and the relative abundances of strains were assigned by the broken-stick model (Supplementary Table S2). Simulated reads were first independently assembled into assemblies by SOAPdenovo2 in each dataset [43], with an empirical k-mer size of 41. Genes were then predicted on assemblies longer than 500bp using MetaGeneMark [42] (default parameters were used except the minimum length of genes was set as 100bp).

Assemblies of each mock dataset were first aligned against the 5 pathogenic *E. coli* reference genomes by BLAT [45]. Those assemblies that have an overall ≥90% overlap and ≥95% identity with the reference genomes were considered as *E. coli* genome-derived (traditional genome alignment-based strategy). Those *E. coli* genome-derived assemblies containing at least three genes (i.e., containing at least two edges) were recruited for construction of a reference pangenome network (RPGN). A query pangenome network (QPGN) was then generated from assemblies selected by the gene alignment-based strategy with *c* = 3 and *r* = 0.5 as described above.

Accuracy of query assembly recruitment was assessed, in respect of conformity and divergence between the RPGN with the QGPN (Supplementary Text 4 and 5 in Supplementary File S2). The result showed that the QPGN recovered 84.3% of node and 84.7% of edge in the RPGN, while falsely included 1.1% of node and 2.2% of edge, which demonstrated the high accuracy of the gene alignment-based strategy for recruitment of metagenome assemblies.

### Construction and visualization of the 760-metagenome pangenome network

Assemblies and representative genes of the 760 metagenomes generated in Reference [24] were used here, since they were produced using identical methods and parameter settings in this study. A pangenome network was generated following steps described above, again using *E. coli* K-12 as the reference, and *c* = 3, *r* = 0.5 for assembly recruitment. The resulting pangenome network was visualized in the same way as visualizing the 5-*E. coli*-genome pangenome network.

### Analysis of subnetworks comprising a pangenome network

10-700 metagenomes were randomly sampled from the above-mentioned 760 metagenomes. For each sub-dataset, a pangenome network was constructed after assembly recruitment using *E. coli* K-12 as the reference genome. For each pangenome network, reference-specific edges were removed before counting the number of subnetworks. Only sub-datasets with a size of recruited assemblies greater than 5 Mb were used to generate the scatterplot, in which a curve with 95% confidence intervals was fitted by the ‘loess’ smoothing method in R [47].

### Computational resources and runtime

Timings for major steps of the MetaPGN pipeline are shown below. Tests were run on a single CPU of an Intel Core Processor (Broadwell) processor with 64 GB of RAM, without otherwise specified. The timings were CPU time including parsing input and writing outputs (h for hours, m for minutes, and s for seconds).

The average time for gene prediction for a mock metagenome was 7s, and it varies depending on the size of a metagenome. The time for redundancy elimination of genes using CD-HIT [44] was 1m 44s for the 5 *E. coli* stains, 50m 19s for the 5 mock datasets. For the 760 metagenomes, to perform redundancy elimination parallelly, we divided all genes into 200 sections, which resulted in 20,101 [N = (*n* + 1) × (*n* ÷ 2) + 1, *n* = 200] clustering tasks, and then submitted each task onto available machines in a high-performance computing cluster. The dividing step took 20m 4s with a peak memory usage of 10GB in the local machine, and the average time for a clustering task was 44m with taking less than 3GB of RAM, consuming total time of 14,814h. The time for recognizing the status (reference-specific, query-specific or shared) for nodes and edges was 10s for the 5 E. coli strains, 1m for the 5 mock datasets and 24m for the 760 metagenomes. Finally, the generation of the pangenome network took less than 1s for the 5 E. coli strains, less than 1s for the 5 mock datasets and 3m 35s for the 760 metagenomes.

### Data availability

Genome sequence of 60 strains (including 5 *E. coli* strains) and the *E. coli* K-12 reference genome were downloaded from the National Center for Biotechnology Information (ftp://ftp.ncbi.nlm.nih.gov/genomes/refseq/bacteria/, Please refer to Supplementary Table S1 for detailed information). Sequencing data of the 760 metagenomes were previously generated in the Metagenomics of the Human Intestinal Tract (MetaHIT) project [21–24], and assemblies of these 760 metagenomes are deposited at EBI under PRJEB28245. The MetaPGN pipeline, related manuals and Cytoscape session files for *E. coli* pangenome networks derived from five pathogenic *E. coli* strains and from 760 metagenomes are available on Github (https://github.com/peng-ye/MetaPGN) and SciCrunch (SCR_016454).

## Additional information

Supplementary Figure S1. Another cluster of genes containing MGEs, flanked by different shared genes on different *E. coli* genomes (manually arranged). Green, blue, red nodes and edges denote reference-specific, shared, and query-specific genes and gene adjacencies, respectively. Triangles represent MGEs. Size of nodes and thickness of edges indicates their weight (occurrence frequency). Numbers alongside shared genes are their indices in the representative gene set, and numbers in parentheses indicate loci of these genes in the reference genome.

Supplementary Figure S2. Examples of arrangement determined by the algorithm. (a) arrangements for shared nodes (blue) and reference-specific nodes (green). (b-e) arrangements for query-specific nodes (red).

Supplementary Table S1. Metadata of isolate genomes used in this study.

Supplementary Table S2. Statistics for the 5 mock metagenomic datasets.

Supplementary Table S3. Tables of nodes and edges in the 5-*E. coli*-genome pangenome network and the 760-metagenome pangenome network.

Supplementary File S1: Texts for, 1) steps for constructing pangenome networks, 2) steps for installing the plug-in and visualizing pangenome networks in Cytoscape.

Supplementary File S2: Texts for, 1) steps for selecting query-specific nodes for arrangement, 2) Comparison of the reference pangenome network (RPGN) and the query pangenome network (RPGN), and 3) detailed definitions of conformity and divergence for nodes and edges.

Supplementary File S3: “5-*E. coli*-genome pangenome network.pdf”, PDF file for *E. coli* pangenome network derived from five pathogenic *E.coli* strains.

Supplementary File S4: “760-metagenome pangenome network.pdf”, PDF file for *E. coli* pangenome network derived from 760 genuine metagenomes.

## Abbreviations

*E. coli*: *Escherichia coli*
LPS: lipopolysaccharide
MGEs: mobile genetic elements
*P. copri*: *Prevotella copri*

## Ethics approval

This study has been approved by the Institutional Review Board on Bioethics and Biosafety (reference number: BGI-IRB 16017).

## Consent for publication

Not applicable.

## Authors’ contributions

J.L. conceived and directed the project. S.T. developed the plug-in. S.T., X.C., Z.Z. and Y.P. developed other codes. Y.P., H.Z., J.L., D.W., S.T. and H.J. performed research. S.T. and Y.P. prepared display items. J.L., H.Z., Y.P., D.W., K.K. and S.T. participated in discussion of the project. Y.P., D.W., H.Z. and S.T. wrote the manuscript. All authors contributed to the revision of the manuscript.

## Acknowledgements

This study was supported by the National Natural Science Foundation of China (No.31601073). We would like to express our appreciation to Dr. Liqiang Li, Dr. Ziqing Deng, Mike Huang-Jingan from BGI-Shenzhen and Prof. Yue Zhang from Sichuan University, for their criticisms and constructive suggestions to this study. We would like to thank Wenchen Song from BGI-Shenzhen for testing the codes. We would also like to extend our gratitude to Chen Ye and Ling Li from BGI-Shenzhen, who made related data publicly available, and Binge Wang and Yanmin Zhao from BGI-Shenzhen for their administrative support.

